# Label-Free Multimodal Volumetric Imaging of Colon Cancer Tissue via Registration of Propagation-Based Phase-Contrast CT, Light-Sheet, and Three-Photon Microscopy

**DOI:** 10.64898/2026.05.21.726767

**Authors:** Christian Dullin, Matthias Schröter, Diana Pinkert-Leetsch, Fernanda Ramos-Gomes, Andrea M. Markus, Jeannine Missbach-Guentner, Hanibal Bohnenberger, Philipp Stroebel, Frauke Alves

## Abstract

Multimodal 3D imaging has emerged as a powerful approach for investigating complex tissue architecture in pathological specimens. Techniques such as propagation-based phase-contrast computed tomography (PCT), light-sheet microscopy (LSM), and three-photon microscopy (3PM) provide complementary information on unlabeled tissue morphology based on distinct intrinsic contrast mechanisms. However, integrating these heterogeneous datasets into a unified spatial framework remains challenging due to differences in imaging geometry, spatial resolution, and modality-specific distortions.

In this study, we present a registration pipeline for spatially aligning volumetric datasets acquired with PCT, LSM, and 3PM from formalin-fixed paraffin-embedded (FFPE) human colon cancer specimens. Biopsies from theses specimens were optically cleared and imaged sequentially using the three high-resolution modalities. To compensate for large positional differences between acquisitions, a three-stage cascade registration strategy was developed, consisting of coarse global alignment on down-sampled data, followed by rigid refinement at intermediate resolution. Mutual information was used as the similarity metric to ensure robust multimodal registration.

The resulting framework enables the generation of spatially aligned multi-channel 3D datasets that combine structural information from X-ray phase-contrast imaging with complementary optical contrast signals. Beyond registration, we demonstrate that the fused six-dimensional feature space can be further exploited for unsupervised tissue characterization using a Gaussian Mixture Model (GMM), enabling data-driven identification of spatially coherent tissue regions without manual annotation.

Qualitative evaluation confirms consistent alignment of major anatomical structures across modalities, while the unsupervised clustering reveals biologically meaningful patterns despite modality-specific noise and resolution differences. While further optimization and validation across larger datasets will enhance its computational efficiency and breadth of application, the approach already demonstrates strong potential for comprehensive tissue analysis and enables scalable, label-free 3D characterization of colon cancer tissue architecture.

## Introduction

Colon cancer is one of the most common malignancies and remains a leading cause of cancer-related mortality worldwide, exhibiting substantial heterogeneity and a complex tumor microenvironment (TME) Sung et al. (2021). Colon cancer can be classified into biologically and clinically distinct subgroups that among other factors show differences in tumor progression and therapeutic response Dunne and Arends (2024). In the era of personalized medicine, structural information obtained by advanced imaging strategies can complement the consensus molecular subtype system based on molecular profiles, enabling a more precise subclassification of colon cancer. Diseases often induce pathological alterations across multiple hierarchical levels, affecting diverse aspects of tissue architecture. Consequently, comprehensive characterization of these changes requires the integration of multiple imaging modalities. Of particular interest are advanced methods such as three-photon microscopy (3PM), light-sheet microscopy (LSM), and propagation-based phase-contrast computed tomography (PCT), which enable volumetric imaging of tissue specimens in three dimensions with near-cellular resolution Yakovlev et al. (2022). These techniques offer significant advantages because they exploit intrinsic tissue properties to generate image contrast, making them highly suitable for label-free imaging approaches.

In contrast, conventional imaging workflows frequently rely on tissue labeling, achieved through diffusion-based staining methods, chemical reactions targeting specific cellular organelles, or the use of labeled antibodies. While effective, these approaches are limited by factors such as incomplete tissue penetration, staining variability, and destructive sample preparation. Label-free imaging overcomes many of these limitations and furthermore enables the investigation of archived tissue specimens, which are commonly preserved as formalin-fixed paraffin-embedded (FFPE) samples Borile et al. (2021). Importantly, the present work builds upon our previous demonstration that FFPE tissues can be reliably imaged using PCT in combination with optical clearing, establishing a robust workflow for volumetric imaging of intact tissue specimens Sagar et al. (2024). When prepared under identical conditions—deparaffinized, rehydrated, optically cleared, and embedded in Phytagel—the resulting datasets acquired by 3PM, LSM, and PCT reveal complementary contrasts arising from the distinct physical principles underlying each modality.

To achieve a comprehensive characterization of tissue specimens, it is essential to integrate these diverse sources of information. In conventional histology and immunohistochemistry (IHC), this integration is typically accomplished by applying different staining protocols to adjacent tissue sections or by using microscopic multiplexing techniques Sun et al. (2025). However, these approaches remain inherently limited to two-dimensional sections and are susceptible to cutting artifacts, tissue deformation, and sampling bias. In recent years, image registration has emerged as a powerful strategy for spatially aligning datasets from different imaging modalities, enabling, for example, the correlation of 2D histological images with 3D PCT data, as demonstrated by Svetlove et al. (2023).

In this study, we present a novel image registration pipeline designed to spatially align volumetric datasets from PCT, three-channel (multiple acquisitions with different filter settings) LSM, and two-channel 3PM into a unified multi-channel representation of human colon cancer tissue. The resulting co-registered dataset combines complementary structural and optical information across modalities and enables comprehensive 3D tissue characterization. Furthermore, we demonstrate that the integrated multimodal volumes can be analyzed using unsupervised learning approaches to identify tissue-specific patterns and morphological features, which are subsequently validated against conventional histology. Our framework therefore establishes a non-destructive and label-free alternative for 3D tissue analysis that avoids physical sectioning, eliminates cutting artifacts, and overcomes the sub-sampling limitations of conventional 2D histology. By preserving the intact tissue architecture, the proposed approach additionally enables quantitative analysis of complex 3D tissue structures and spatial relationships within the TME of human colon cancer specimens.

## Results

### Workflow

The major processing steps of the registration pipeline are summarized in Figure 1. Deparaffinized core punch biopsies (6 mm diameter) obtained from FFPE human colon cancer specimens were embedded in Phytagel and optically cleared according to the protocol described in Sagar et al. (2024). This preparation procedure was previously established for both PCT and LSM imaging and was additionally applied for 3PM acquisition. In this presented study, while the acquisition order of PCT and LSM varied between specimens, 3PM imaging was consistently performed last. As a result, three volumetric image datasets were generated for each specimen, capturing either the complete tissue volume or a substantial fraction thereof (Figure 1).

**Figure 1.**
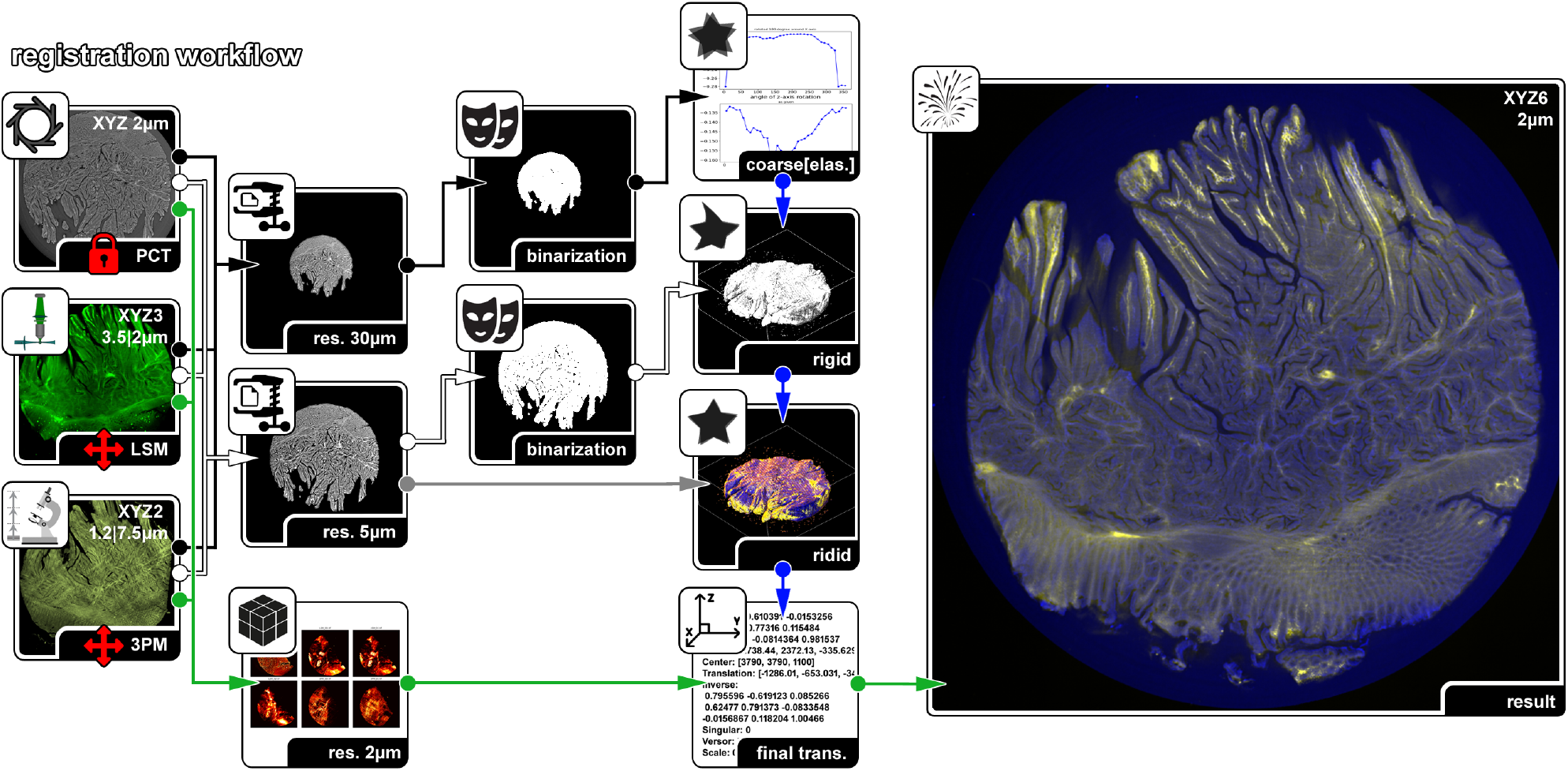
Registration workflow. The three input data sets - PCT, LSM, and 3PM are resampled to two sets with isotropic voxel sizes of 30 µm and 5 µm, respectively. PCT is treated as static reference, while LSM and 3PM are adapted to match, using always the respected first channel of the data sets. The low resolution binarized data was used for coarse match testing rotation angles in 5°increments and potential flip. The found transformation was then applied to the 5 µm binarized data and further refined by using the non-binarized 5 µm in a final step. The resulting transformation was then applied to all six channels resampled to 2 µm isotropic voxel size. The process was conducted for the PCT-LSM and PCT-3PM pairs separately. PCT = Propagation-based phase-contrast computed tomography. LSM = Light-sheet microscopy. 3PM = Three-photon microscopy. Arrow colors indicate the different processing streams: black arrows denote the 30 µm low-resolution data used for coarse registration, white arrows the binarized 5 µm data used for refined alignment, gray arrows the non-binarized 5 µm data used for final refinement, blue arrows the sequential transformation steps, and green arrows the application of the final transformations to the high-resolution 2 µm multichannel data.

The PCT datasets consisted of a single intensity channel with an isotropic voxel size of 2 µm. The LSM datasets provided three autofluorescence channels acquired using excitation/emission filter combinations of 470/525 nm, 560/620 nm, and 710/810 nm, respectively, with an in-plane resolution of approximately 3.5– 4 µm and a slice thickness of 2 µm. The 3PM datasets comprised two channels acquired at an excitation wavelength of 1300 nm, corresponding to forward-scattered second- and third-harmonic generation signals detected using 647LP and 432/36 filter configurations, respectively. The resulting datasets exhibited an in-plane resolution of 1.2 µm and a slice thickness of 7.5 µm.

For multimodal image registration, two principal aspects had to be addressed: (i) the expected geometric transformations between datasets and (ii) the metric used to evaluate spatial correspondence. The deformation model needed to account for substantial positional and rotational differences arising from specimen tilting and flipping during transport and PCT acquisition. Furthermore, small inaccuracies in the nominal voxel sizes of the different imaging modalities required the inclusion of a limited scaling component within the transformation model.

In addition, the LSM datasets exhibited modality-specific optical distortions caused by residual scattering within the optically cleared tissue. Despite clearing, scattering effects resulted in a slight broadening of the light sheet during propagation through the specimen. In combination with specimen rotation during volumetric acquisition, this produced a characteristic biconcave and rotationally symmetric deformation pattern within the reconstructed volumes.

As these effects could not be robustly captured within a single optimization step, a cascaded multi-stage registration strategy was employed:

- Initial estimation of large-scale translation, rotation, flipping, and scaling using heavily downsampled and binarized datasets at an isotropic resolution of 30 µm.
- Refinement of the global transformation parameters using binarized datasets resampled to an isotropic voxel size of 5 µm.
- Intensity-based refinement at 5 µm isotropic resolution using the original grayscale information of the datasets.
- Application of the optimized transformations to all modalities resampled to an isotropic voxel size of 2 µm, corresponding to the native resolution of the PCT reference dataset.

Due to the substantial differences in image contrast and signal characteristics between modalities, mutual information was selected as the similarity metric, specifically employing Mattes mutual information as described in Mattes et al. (2003). PCT datasets, which typically provided isotropic and minimally distorted volumetric information, served as the fixed reference images, whereas the LSM and 3PM datasets were treated as moving images. Since the multi-channel LSM acquisitions were performed sequentially without repositioning of the specimen, the transformation parameters were estimated using the first channel and subsequently applied to the remaining two channels. The same procedure was used for the two-channel 3PM datasets. Registration of LSM and 3PM data was performed independently, each using the corresponding PCT dataset as reference.

To further compensate for residual specimen deformations caused by mounting conditions and temperature variations between imaging modalities, an optional elastic registration step was introduced following the coarse alignment at 30 µm resolution. However, this non-rigid refinement was only applicable when the complete specimen surface was visible in both image pairs under consideration (PCT–LSM or PCT–3PM).

The final outcome of the workflow was a co-registered six-channel volumetric dataset with an isotropic voxel size of 2 µm for each specimen.

### Registration result

Figure 2 demonstrates the successful alignment of the three imaging modalities. Despite the fact that each modality emphasizes different tissue properties and structural features, the corresponding anatomical regions are consistently co-registered across all datasets. This is particularly notable given that parts of the specimen are absent in the 3PM acquisition (lower right corner, labeled & in Figure 2), illustrating the robustness of the registration pipeline against incomplete image content.

**Figure 2.**
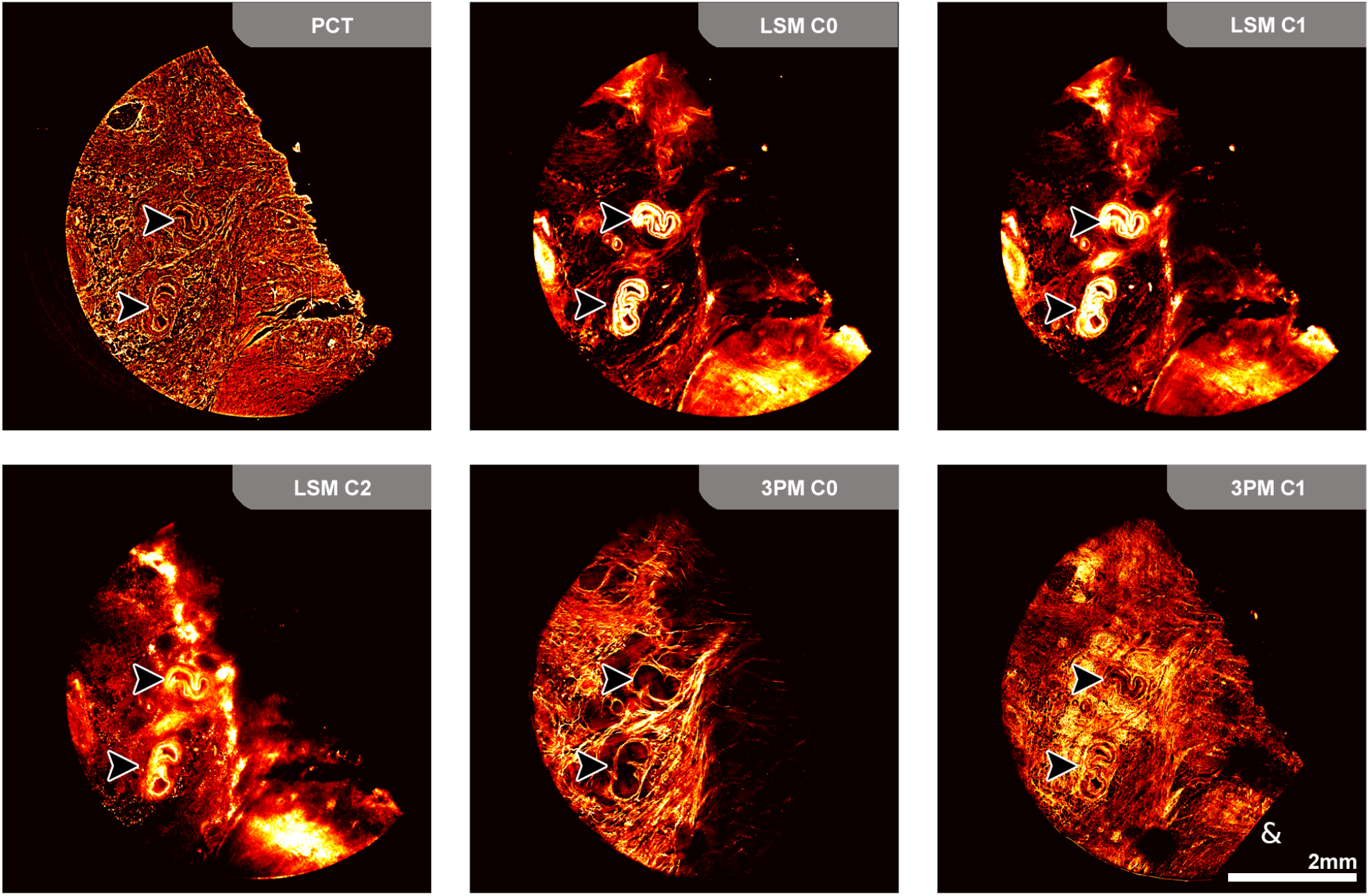
Fusion results. The 2D images depict corresponding slices from the three imaging modalities and their respective filter channels (C0, C1, C2) following resampling to an isotropic voxel size of 2 µm and multimodal registration. The successful spatial alignment is demonstrated by the consistent positioning and orientation of anatomical structures across modalities, as exemplified by the matching vessel cross-sections visible in all datasets (arrow heads). PCT = Propagation-based phase-contrast computed tomography. LSM = Light-sheet microscopy. 3PM = Three-photon microscopy.

The accurate spatial alignment enabled the generation of a fused 3D red–green–blue (RGB) representation of the multimodal data (Figure 3). For visualization, each channel was individually normalized according to:

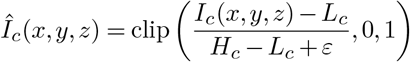

where:

**Figure 3.**
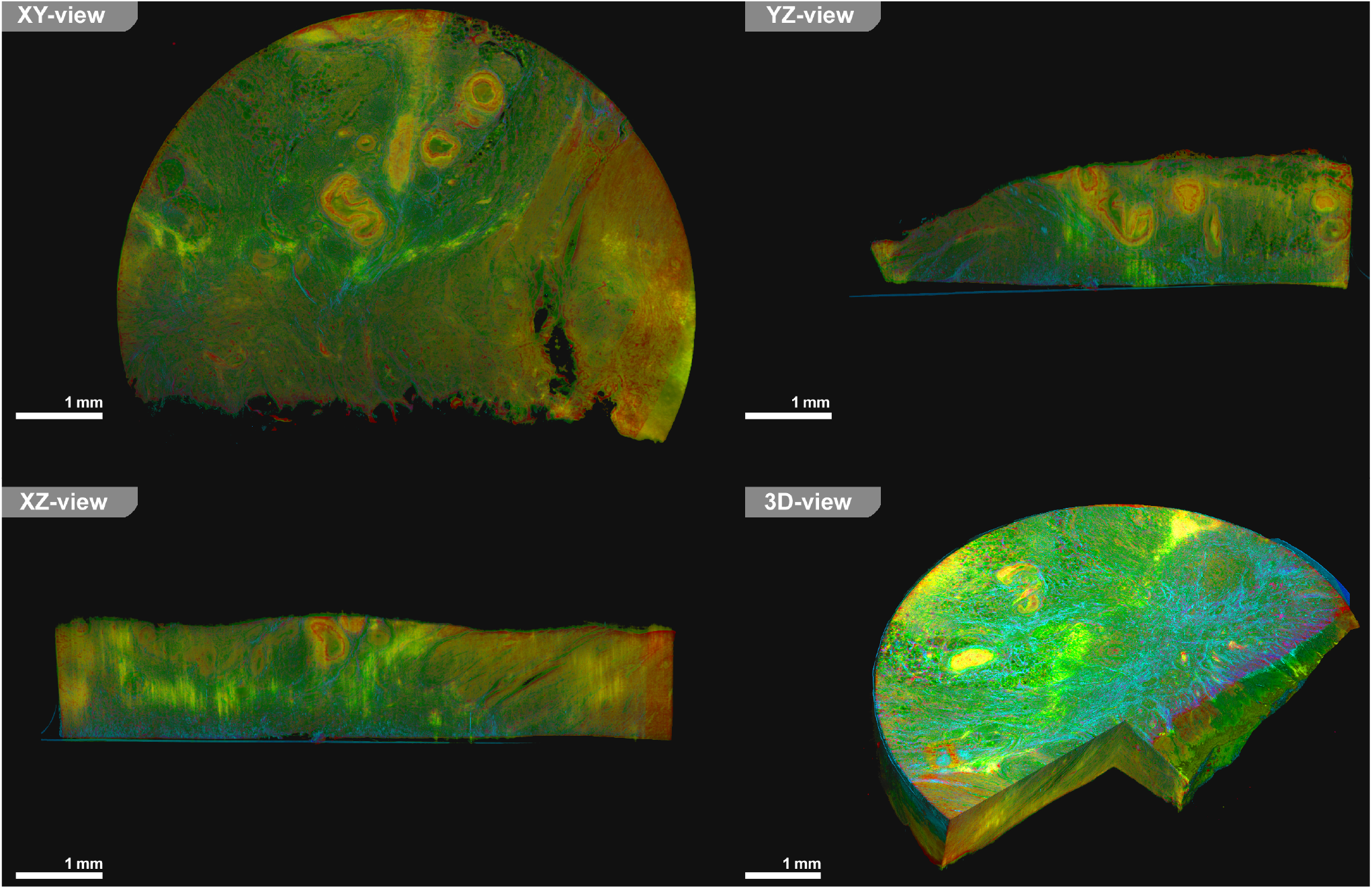
Color-merged representation. demonstrates that different tissue structures exhibit distinct composite color signatures reflecting modality-specific contrast contributions. However, especially in the XZ and YZ views, the limited axial resolution and slice thickness of LSM and 3PM become apparent, resulting in a noticeable blurring and elongation of structures along the z-direction. In contrast, the 3D rendering emphasizes that the dataset represents a true volumetric specimen, which can be virtually resliced in arbitrary positions and orientations without loss of spatial consistency, highlighting the advantage of isotropic reconstruction for 3D tissue analysis. LSM = Light-sheet microscopy. 3PM = Three-photon microscopy.

- *I*_*c*_ denotes image channel *c* ordered as follows: (PCT, LSM470*/*525nm, LSM560*/*620nm, LSM710*/*810 nm, 3PM1300 nm, 3PM1650 nm),
- *L*_*c*_ and *H*_*c*_ represent the lower and upper intensity normalization bounds, respectively,
- *ε* is a small stabilization constant preventing division by zero.

The normalized channels were subsequently combined into an RGB representation according to:

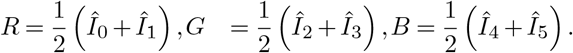

This color fusion facilitates simultaneous visualization of complementary structural information provided by the different imaging modalities and highlights the spatial relationships between tissue components within the intact 3D specimen. However, despite its utility for qualitative assessment, the color-merged representation does not readily allow for the reliable identification or characterization of distinct tissue types, as overlapping intensity contributions and complex multimodal contrast mechanisms obscure clear separation of underlying tissue classes (Figure 3).

### Tissue characterization by Gaussian Mixture Model estimation

Assuming accurate spatial alignment across all three imaging modalities, a central question is which additional biological information can be extracted from the resulting multimodal dataset. In the absence of ground-truth annotations for specific tissue or cell types, this problem can be formulated as an unsupervised learning task in a six-dimensional feature space defined by the registered imaging channels. The underlying hypothesis is that distinct tissue and cellular states are characterized by unique, modality-dependent intensity signatures across the combined PCT, LSM, and 3PM channels.

In practice, however, the feature space is expected to exhibit substantial overlap between classes. This is primarily due to residual registration inaccuracies, differences in voxel size between modalities, the strongly non-isotropic sampling characteristics of LSM and 3PM, deformation effects introduced during specimen handling, and the limited spatial resolution at tissue interfaces. In addition, missing signal regions in the 3PM data and reconstruction artifacts in LSM (e.g., signal voids or halos outside the specimen boundary) further complicate the structure of the feature distribution.

To account for this complexity, we model the joint intensity distribution using a Gaussian Mixture Model (GMM), which assumes that the observed data are generated by a weighted sum of multivariate Gaussian components. The model can be written as:

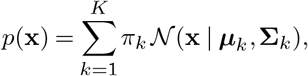

where **x** ∈ ℝ^6^ denotes the six-dimensional feature vector at each voxel, *π*_*k*_ are the mixture weights with 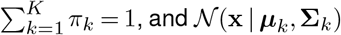 denotes a multivariate Gaussian distribution with mean vector ***µ***_*k*_ and covariance matrix **Σ**_*k*_.

The optimal number of components *K* was estimated using the Bayesian Information Criterion (BIC) computed on a representative subset of the data. This procedure resulted in *K* = 28 components. Importantly, this number should not be interpreted as corresponding directly to distinct biological tissue or cell types. Instead, the learned components capture a mixture of biological variability, imaging modality-specific effects, missing data regions (e.g., absent 3PM signal), and acquisition-related artifacts, including background and boundary effects observed in the LSM data.

Despite this overcomplete representation, the GMM provides a useful probabilistic partitioning of the high-dimensional intensity space and serves as a basis for subsequent tissue characterization and spatial interpretation of multimodal signal distributions.

The GMM-based tissue characterization (Figure 4) further revealed spatially coherent regions that corresponded to major histological structures, despite the absence of supervision. While larger anatomical features such as adipose tissue (black arrow head) and vascular components (white arrow head) were consistently identified across modalities, finer structures, particularly in the colonic epithelium (§), showed increased cluster ambiguity due to limited axial resolution and partial volume effects in LSM and 3PM. These limitations manifested as mixed or fragmented component assignments rather than well-defined discrete clusters.

**Figure 4.**
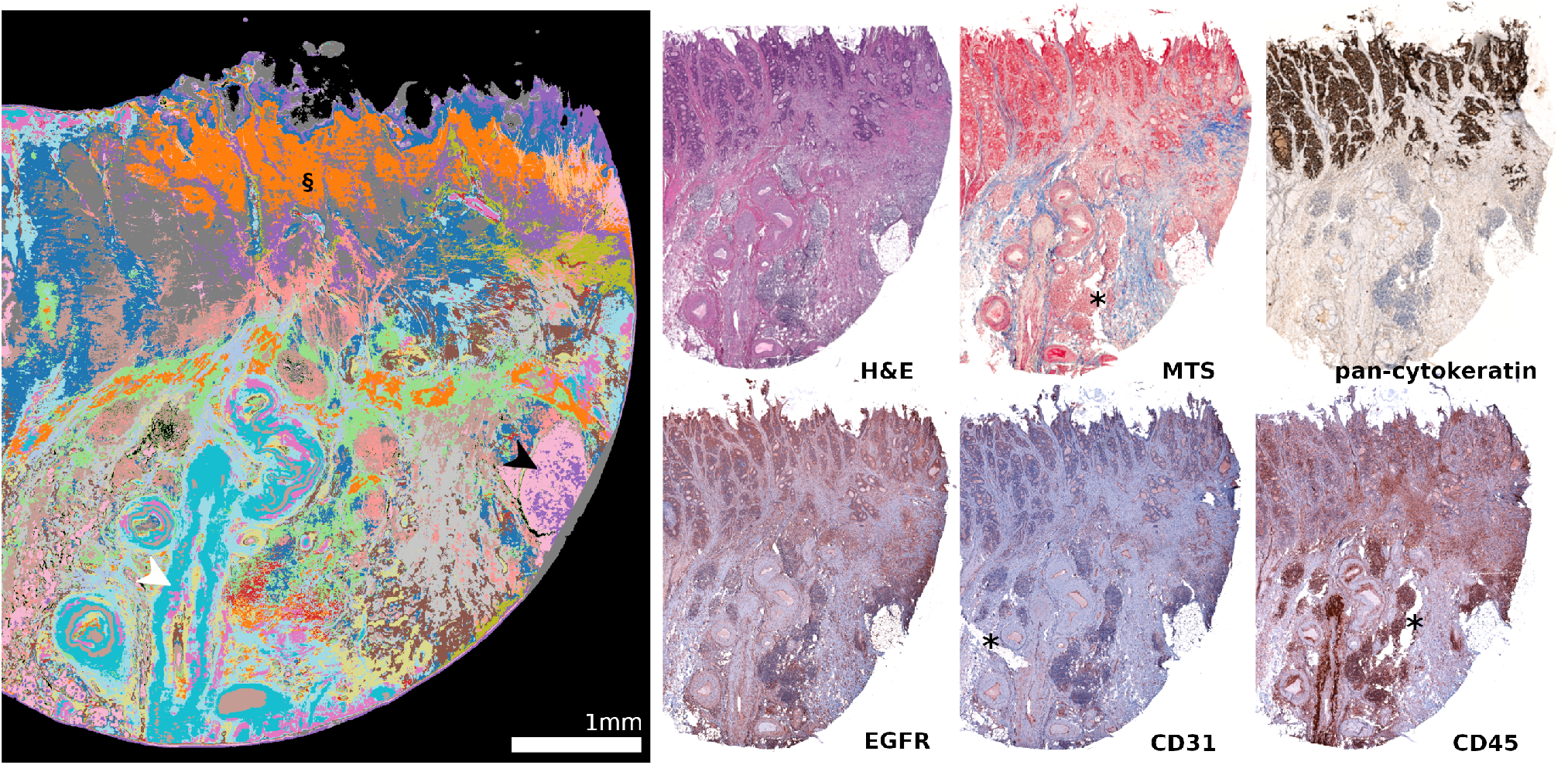
The GMM tissue classification result. (left panel) shows a pseudo-color representation of voxel-wise classification of a colon cancer dataset using the 28-component GMM. Peripheral light-gray regions indicate areas of poor data correspondence between modalities. Distinct components can be associated with specific tissue structures, e.g. pink (black arrowhead) corresponds to fatty tissue and turquoise (white arrowhead) to the vessel wall. In regions with fine-scale tissue structures (§), the limited axial sampling of LSM and 3PM leads to heterogeneous cluster assignments. The right panel shows corresponding histological sections, which are subject to sectioning artifacts (*) and do not allow repeated staining of the same slice. GMM = Gaussian Mixture Model. H&E = Hematoxylin and eosin. MTS = Masson’s trichrome staining. EGFR = Epidermal growth factor receptor. CD31 = Platelet endothelial cell adhesion molecule-1. CD45 = Protein tyrosine phosphatase receptor type C.

Correlative histological analysis was performed on adjacent 2 µm sections obtained from the central region of the 6 mm tissue biopsy using hematoxylin and eosin (H&E) for general cell morphology, Masson’s trichrome staining (MTS) for collagen/connective tissue, and IHC labeling against the tumor markers pan-cytokeratin and EGFR (epidermal growth factor receptor), as well as CD31 (platelet endothelial cell adhesion molecule-1), and CD45 (protein tyrosine phosphatase receptor type C) to highlight endothelial and immune cell populations. In contrast to classical histology, which provides high lateral resolution but is limited to destructively obtained single sections that are susceptible to sectioning artifacts (*) and prevent repeated staining of the identical slice, the proposed volumetric approach preserves the 3D integrity of the tissue and enables a more comprehensive interpretation of structural context.

## Discussion

A central challenge in multimodal imaging of biological tissue is the reliable spatial integration of datasets acquired with fundamentally different imaging principles, spatial resolutions, and deformation characteristics. In this study, we present a cascade-based registration framework for aligning PCT, LSM, and 3PM datasets acquired from the same 6 mm core biopsy of FFPE colon cancer specimens. The results demonstrate that, despite substantial differences in contrast mechanisms and acquisition geometry, a mutual information–based registration strategy, combined with a coarse-to-fine transformation cascade, enables robust alignment of major anatomical structures across modalities.

Using PCT as an isotropic structural reference is particularly beneficial, as it provides a geometrically stable backbone onto which the more deformation-prone optical modalities can be mapped. The initial coarse alignment resolves large rigid transformations arising from specimen handling, while subsequent refinement stages compensate for smaller-scale deformations, particularly those observed in LSM data. The final fused representation integrates complementary contrast information from all modalities into a coherent volumetric dataset, enabling more comprehensive characterization of tissue architecture than any individual modality alone.

Compared with classical histology, which relies on destructive 2D sectioning and is therefore prone to cutting artifacts, tissue deformation, and sampling bias, the proposed framework preserves the volumetric integrity of the specimen. This enables arbitrary virtual sectioning and consistent spatial interrogation across the entire tissue volume. Similar advantages of volumetric imaging for the spatially resolved analysis of heterogeneous pathological structures have been demonstrated in prior multimodal imaging studies Benke et al. (2025).

In this sense, the approach moves towards a form of “continuous histology” in three dimensions, thereby extending approaches to integrate 2D histology into PCT for instance to provide specificity for segmentation approaches, such as demonstrated by Reiser et al. for segmenting the axial connective fiber architecture in lung tissue Reiser et al. (2024).

Related approaches for reconstructing 3D tissue representations from histological or micro-computed tomography (microCT) data have primarily focused on 2D-to-3D registration of serial sections or alignment of histology to PCT volumes Pichat et al. (2018), as demon-strated in previous work on multimodal histology reconstruction Song et al. (2013) and CT-to-histology registration frameworks Albers et al. (2021). In contrast, the present work extends these concepts by jointly integrating multiple high-resolution 3D modalities with fundamentally different contrast mechanisms, rather than relying on 2D section stacks or single-modality reconstruction.

A key outcome of this work is that the resulting multimodal dataset can serve as a basis for exploratory analyses such as feature correlation and unsupervised learning. In this context, early results suggest that biologically meaningful structures may emerge from the joint intensity space even without explicit annotations. This motivates a link to emerging “virtual staining” approaches, where phase-contrast or label-free volumetric imaging is mapped to histology-like contrast using learning-based transformations Almagro-Pérez et al. (2025). However, unlike purely generative histology translation methods, our framework remains physically grounded in multimodal acquisition rather than synthetic contrast generation, which reduces the risk of hallucinated structures but also limits direct interpretability in histology terms.

The three imaging modalities were selected to provide complementary contrast across different spatial scales and tissue properties within the optically cleared FFPE colon carcinoma specimens. Propagation-based phase-contrast computed tomography (PCT) at the synchrotron provided a label-free, volumetric overview of the entire biopsy core at 2 µm isotropic voxel size, exploiting subtle differences in electron density and refractive index between tissue compartments—such as glandular epithelium, stroma, vascular structures, and necrotic regions—without requiring any staining Svetlove et al. (2023). Light-sheet microscopy (LSM) captured mesoscale autofluorescence across three spectral channels, where the 470/525 nm channel predominantly reflects NADH and structural protein emission, the 560/620 nm channel captures FAD and collagen-associated fluorescence, and the 710/810 nm channel isolates longer-wavelength autofluorescent species. Together, these channels enabled discrimination of metabolically and structurally distinct tissue regions throughout the cleared specimen volume Jahr et al. (2015); Deal et al. (2018). Three-photon microscopy (3PM) provided high-resolution, label-free structural contrast through second- and third-harmonic generation. SHG at 1300 nm excitation selectively highlighted fibrillar collagen within the tumor stroma due to its noncentrosymmetric organization, enabling visualization of stromal remodeling and extracellular matrix organization, while THG at 1650 nm marked interfaces between optically heterogeneous compartments such as lipid-rich membranes, cell boundaries, glandular interfaces, and extracellular matrix structures, making it particularly sensitive to tissue architecture at the cellular scale Debarre et al. (2006); Rehberg et al. (2011). Together, these modalities span from millimeter-scale tissue organization down to subcellular structural features, enabling multiscale characterization of tumor morphology, stromal composition, and intrinsic tissue heterogeneity without exogenous labeling.

Importantly, the combination of PCT, LSM, and 3PM—each providing tissue contrast through distinct intrinsic physical mechanisms—may enable visualization of structural features that remain inaccessible to conventional histological workflows. Unlike classical histopathology, which relies on predefined molecular stains or antibody-based labeling, this multimodal label-free approach exploits endogenous optical and structural contrast inherent to the tissue itself. As a result, it has the potential to reveal morphologies, tissue interfaces, and microarchitectural patterns for which no dedicated staining protocol currently exists or is routinely applied in clinical pathology. In this way, multimodal label-free imaging may extend beyond the limitations of conventional histology by accessing complementary contrast dimensions across multiple spatial scales. While combinations of PCT with other modalities, including optical microscopy Takeda et al. (2000), SEM Reiser et al. (2024), and AFM D’Amico et al. (2024), have previously been reported, to our knowledge the integrated application of PCT, LSM, and 3PM for the characterization of human colon carcinoma specimens has not yet been demonstrated.

Nevertheless, interpretability remains limited in regions containing fine tissue structures, where resolution constraints, partial volume effects, and modality-specific blur reduce cluster separability. Moreover, the unsupervised clustering approach (e.g., GMMs) does not inherently guarantee correspondence to biologically discrete cell types, but rather identifies statistically separable intensity regimes, which may reflect a mixture of tissue composition, imaging artifacts, and acquisition-specific noise.

An important practical constraint of the presented pipeline is that integration of PCT, LSM, and 3PM currently requires optically cleared specimens. While PCT imaging alone can in principle also be performed on non-cleared, paraffin-embedded tissue Reiser et al. (2024); Albers et al. (2021), the optical clearing required for LSM and 3PM introduces an additional source of tissue alteration. In particular, dehydration and solvent-based clearing (e.g., benzyl alcohol/benzyl benzoate or ethyl cinnamate) may induce tissue shrinkage, swelling, or non-linear deformations, which can differ between samples and spatial locations. These effects can complicate inter-modality registration, especially when comparing modalities acquired before and after clearing, and may represent a non-negligible source of systematic distortion in the combined dataset.

Several additional practical and methodological limitations should be considered. The registration strategy assumes that the complete or nearly complete outer geometry of the specimen is visible in all modalities, which requires full-volume acquisition even when only subregions are of interest. This increases acquisition time and data size, particularly for 3PM, where mosaic imaging is required to cover larger fields of view. In addition, all datasets are resampled to a common isotropic resolution, which inevitably reduces the effective resolution of higher-resolution modalities and introduces interpolation artifacts, especially in strongly anisotropic LSM data.

Another limitation is the lack of explicit modeling of modality-specific imaging physics during registration. While mutual information provides robustness to intensity differences, it does not explicitly account for scattering-induced blur in LSM, harmonic generation signal bias in 3PM, or reconstruction artifacts in PCT. This may introduce subtle systematic misalignments in regions with weak structural correspondence.

Finally, the computational cost of the full pipeline remains significant, requiring several hours per dataset on high-memory workstations. As a result, scalability to large cohorts remains limited without further optimization or GPU-accelerated registration strategies.

Future work should therefore focus on three main directions: (i) incorporation of physics-aware or learning-based registration models that explicitly account for modality-specific distortions, (ii) validation of biological interpretability using targeted histology and IHC as ground truth, and (iii) scaling the framework towards cohort-level analysis. In particular, systematic comparison between unsupervised multimodal classification and classical histopathological grading will be essential to assess clinical relevance.

Ultimately, multimodal volumetric integration combined with unsupervised tissue characterization represents a step towards non-destructive, label-free 3D histology. However, careful validation against conventional histology remains essential before such approaches can replace or augment established diagnostic workflows.

## Methods

### Sample preparation

Cylindrical biopsies were obtained from FFPE human colon carcinoma specimens using a 6-mm skin punch biopsy needle (pfm medical, Germany). Samples were first incubated at 60°C for 30 minutes in a heating cabinet, followed by incubation in xylene for 4 hours at 60°C under gentle agitation. Subsequently, samples were processed in a tissue infiltration system: 59 hours in xylene at room temperature (RT), followed by two steps in 100% EtOH for 12 hours at RT, two steps in 96% EtOH for 4 hours, two steps in 75% EtOH for 4 hours, and single incubations in 60% and 50% EtOH for 4 hours each. Finally, samples were rehydrated in water for 4 hours at RT.

The specimens were then embedded in 0.7% Phytagel (Sigma-Aldrich, USA) prepared in water, using cylindrical molds (e.g., 12 mm syringe). Embedding was performed at 4°C for 30 minutes. Embedded samples were transferred to phosphate-buffered saline (PBS) and gently rocked for 60 minutes at RT. Dehydration was then repeated in graded ethanol series (30% for 4 hours, 50% overnight, 70% for 4 hours, 90% for 4 hours, and 100% for 4 hours, all at RT). Finally, samples were cleared in a 1:1 mixture of benzyl alcohol and benzyl benzoate (BABB; Sigma-Aldrich, USA) at RT in the dark and stored in 5 ml tubes filled with BABB.

### Light-sheet microscopy (LSM)

Light-sheet microscopy was performed in ethyl cinnamate (ECI) using a Blaze system (Miltenyi Biotec, Germany) equipped with a 1x objective and 1.66x zoom. The light sheet was generated with a numerical aperture (NA) of 0.163 and a thickness of 3.9 µm, with a sheet width of 70%, and a z-step size of 2 µm.

Imaging was performed using excitation/emission filter sets of 470/525 nm, 560/620 nm, and 710/810 nm, resulting in an anisotropic voxel size of approximately 3.5 µm *×* 3.5 µm *×* 2.0 µm.

### Three-photon microscopy (3PM)

For 3PM, samples were removed from BABB and excess Phytagel was carefully removed. Specimens were mounted on low-adhesion glass-bottom dishes (35 mm; ibidi, Germany), covered with a 25 mm coverslip, and sealed with vaseline (Molyduval, Germany).

Label-free second and third-harmonic generation (SHG/THG) imaging was performed using an upright TriM Scope Matrix multiphoton microscope (Miltenyi Biotec, Germany) equipped with a tunable femtosecond laser source (1300–1700 nm; CRONUS-3P, Light Conversion, Lithuania). System alignment and illumination settings were verified prior to acquisition using a calibration slide (Chroma, USA).

Acquisition parameters included a 25x Olympus water-immersion objective and oil-immersion condenser, excitation wavelengths of 1300 nm and 1650 nm, polarization angle of 0°, field of view of approximately 300 µm, pixel sizes of 600–1100 nm, and z-step sizes of 10– 20 µm. Mosaic acquisitions with 10% overlap were used when necessary. After imaging, samples were returned to BABB.

### Propagation-based phase-contrast computed tomography (PCT)

Specimens were scanned in sealed 5 ml tubes filled with BABB at the SYRMEP beamline (SYnchrotron Radiation for MEdical Physics) of the Elettra synchrotron in Trieste, Italy. The beamline was operated in pink beam mode Dullin et al. (2021), with a sample-to-detector distance of 50 cm to enhance phase contrast for weakly absorbing tissues.

Full 360°rotation scans were acquired with a voxel size of 2 µm, resulting in a field of view of approximately 7.5*×* 7.5*×* 4 mm^3^. A total of 3600 projections were recorded with exposure times of 20–50 ms at beam energies of 2.0–2.4 GeV, yielding total scan times of 1.2– 3 minutes. For specimens exceeding 4 mm in height, vertically overlapping scans were acquired.

Phase retrieval was performed using a single-distance Paganin filter (TIE_HOM approach Paganin et al. (2002)) with a delta-to-beta ratio of 100, followed by filtered back-projection reconstruction using the SYRMEP Tomo Project (STP) Brun et al. (2015).

### Histology

Following completion of multimodal imaging, the specimens were removed from the Phytagel matrix and thoroughly washed to eliminate residual clearing agent (BABB). The samples were subsequently re-embedded in paraffin using standard dehydration to ensure compatibility with microtome sectioning Sagar et al. (2024). From each specimen, serial sections with a thickness of 2 µm were obtained from the central region of the tissue biopsy to ensure spatial correspondence with the volumetric imaging data. Sections were cut using a rotary microtome and mounted on glass slides for histological and IHC analysis on adjacent sections.

Conventional histological staining was performed on tissue sections using H&E to assess overall tissue morphology, as well as MTS (Sigma–Aldrich, USA) to visualize collagen-rich extracellular matrix structures, according to the manufacturer’s instructions. In addition, IHC staining was performed. For cytokeratin staining, which was used to identify tumors of epithelial origin, tissue sections were incubated in EnVision Flex target retrieval solution (Dako, USA) at high pH, followed by incubation with a primary monoclonal mouse anti-human cytokeratin antibody (ready-to-use, Dako, USA, GA053) for 12.5 min at RT, and subsequently with the secondary antibody system (EnVision Flex+, Dako, USA). Mayer’s hematoxylin was used for counterstaining. Additional IHC staining was performed on adjacent sections following pre-treatment with target retrieval solution (pH 9, 100°C, 20 min), H_2_O_2_ (10 min, RT), and Sea Block (20 min, RT). The following primary antibodies were applied to assess vascularization, immune cell infiltration, epithelial tumor origin, and tumor-associated receptor expression: CD31 (polyclonal rabbit antibody, ab28364, Abcam, UK, 1:50; endothelial marker), CD45 (rabbit monoclonal antibody, ab208022, Abcam, 1:4000; pan-leukocyte marker), and EGFR (mouse monoclonal antibody, MA5-16359, Thermo Fisher Scientific, USA, 1:100). Sections were incubated overnight with the respective primary antibodies on adjacent tissue sections, followed by 30 min incubation at RT with horseradish peroxidase (HRP)-conjugated goat anti-rabbit or antimouse secondary antibodies (Histofine, Nichirei Biosciences, Japan). AEC (3-amino-9-ethylcarbazole) substrate was used as chromogen, and sections were briefly counterstained with hematoxylin (Hämalaun) for 3 s, followed by a 5 min bluing step in running tap water. Finally, sections were mounted using Aquatex (Sigma– Aldrich, USA).

Stained sections were imaged using the Carl Zeiss AG Axio Scan.Z1 slide scanner (Carl Zeiss AG, Germeny) at high resolution and subsequently used for qualitative comparison and validation of structural features observed in the multimodal 3D imaging datasets.

### Software

Image registration was performed using SimpleITK Lowekamp et al. (2013) in Python. PCT reconstruction and phase retrieval were carried out using the SYRMEP Tomo Project (STP v1.6) Brun et al. (2015). Fiji v1.54 Schindelin et al. (2012) was used for basic image processing. The 3D visualization and rendering were performed with VGStudio Max v3.1 (Volume Graphics, Germany). Figures were prepared using Adobe Photoshop (Adobe Inc., USA)

### Ethics

Human colon carcinoma specimens were collected from patients at the Institute of Pathology, University Medical Center Göttingen (Germany), where routine diagnostic procedures were performed. The study was approved by the local ethics committee of the University Medical Center Göttingen (approval number 24/4/20) and conducted in accordance with the Declaration of Helsinki.

3PM: Three-photon microscopy
AEC: 3-amino-9-ethylcarbazole
AFM: Atomic force microscopy
BABB: Benzyl alcohol/benzyl benzoate
BIC: Bayesian Information Criterion
CD31: Platelet endothelial cell adhesion molecule-1
CD45: Protein tyrosine phosphatase receptor type C
ECI: Ethyl cinnamate
EGFR: Epidermal growth factor receptor
FAD: Flavin Adenine Dinucleotide
FFPE: Formalin-fixed paraffin-embedded
GMM: Gaussian Mixture Model
HE: Hematoxylin and eosin
HRP: Horseradish peroxidase
IHC: Immunohistochemical
LSM: Light-sheet microscopy
microCT: High-resolution micro-computed tomography
MTS: Masson’s trichrome staining
NA: Numerical aperture
NADH: Nicotinamide Adenine Dinucleotide (Reduced Form)
PBS: Phosphate-buffered saline
PCT: Propagation-based phase-contrast computed tomography
RGB: Red–green–blue
RT: Room temperature
SEM: Scanning electron microscopy
SHG: Second-harmonic generation
STP: SYRMEP Tomo Project
THG: Third-harmonic generation
TME: Tumor microenvironment

## Acknowledgements

We thank Sabine Wolfgram, Sarah Garbode, Stefan Lesemann, Bettina Jeep, Julia Fascher (UMG), as well as Bärbel Heidrich and Regine Kruse (MPI-NAT), for their excellent technical assistance with sample preparation, histological and immunohistochemical staining, light-sheet image acquisition (J.F.), and data reconstruction (S.L.). We further gratefully acknowledge the SYRMEP team for their continuous technical support throughout the project.

## Bibliography

Albers, J., Svetlove, A., Alves, J., Kraupner, A., di Lillo, F., Markus, M. A., Tromba, G., Alves, F., and Dullin, C. Elastic transformation of histological slices allows precise co-registration with microCT data sets for a refined virtual histology approach. Scientific Reports, 11(1): 10846, 2021.

Almagro-Pérez, C., Peruzzi, N., Galambos, C., Song, A. H., Brunnström, H., Gawlik, K. I., Stampanoni, M., Tran-Lundmark, K., and Lovric, G. Histology-guided 3D virtual staining of microCT-imaged lung tissue via deep learning. bioRxiv, pages 2025–10, 2025.

Benke, C. V., Duerr, J., Engel, A., Dullin, C., Stiller, W., Horstmann, H., Redenbach, C., Ackermann, M., Kauczor, H. U., Kuner, T., Wielputz, M. O., Mall, M. A., and Wagner, W. L. High-resolution multimodal imaging reveals spatial and temporal heterogeneity of airway mucus plugging in mice with muco-obstructive lung disease. Sci Rep, 15(1): 41760, 2025. doi: 10.1038/s41598-025-22537-7.

Borile, G., Sandrin, D., Filippi, A., Anderson, K. I., and Romanato, F. Label-free multiphoton microscopy: much more than fancy images. International Journal of Molecular Sciences, 22(5):2657, 2021.

Brun, F., Pacilè, S., Accardo, A., Kourousias, G., Dreossi, D., Mancini, L., Tromba, G., Pugliese, R., et al. Enhanced and flexible software tools for X-ray computed tomography at the Italian synchrotron radiation facility Elettra. Fundamenta Informaticae, 141 (2-3):233–243, 2015.

Deal, J., Harris, B., Martin, W., Lall, M., Lopez, C., Rider, P., Boudreaux, C., Rich, T., and Leavesley, S. J. Demystifying autofluorescence with excitation scanning hyperspectral imaging. In Imaging, Manipulation, and Analysis of Biomolecules, Cells, and Tissues XVI, volume 10497, pages 129–136. SPIE, 2018.

Debarre, D., Pena, A.-M., Supatto, W., Boulesteix, T., Strupler, M., Sauviat, M.-P., Martin, J.-L., Schanne-Klein, M.-C., and Beaurepaire, E. Second-and third-harmonic generation microscopies for the structural imaging of intact tissues. Medecine sciences: M/S, 22 (10):845–850, 2006.

Dullin, C., di Lillo, F., Svetlove, A., Albers, J., Wagner, W., Markus, A., Sodini, N., Dreossi, D., Alves, F., and Tromba, G. Multiscale biomedical imaging at the SYRMEP beamline of Elettra-closing the gap between preclinical research and patient applications. Physics Open, 6:100050, 2021.

Dunne, P. D. and Arends, M. J. Molecular pathological classification of colorectal cancer—an update. Virchows Archiv, 484(2):273–285, 2024.

D’Amico, L., Svetlove, A., Longo, E., Meyer, R., Senigagliesi, B., Saccomano, G., Nolte, P., Wagner, W. L., Wielpütz, M. O., Leitz, D. H., et al. Characterization of transient and progressive pulmonary fibrosis by spatially correlated phase contrast microct, classical histopathology and atomic force microscopy. Computers in Biology and Medicine, 169: 107947, 2024.

Jahr, W., Schmid, B., Schmied, C., Fahrbach, F. O., and Huisken, J. Hyperspectral light sheet microscopy. Nature communications, 6(1):7990, 2015.

Lowekamp, B. C., Chen, D. T., Ibáñez, L., and Blezek, D. The design of SimpleITK. Frontiers in neuroinformatics, 7:45, 2013.

Mattes, D., Haynor, D. R., Vesselle, H., Lewellen, T. K., and Eubank, W. PET-CT image registration in the chest using free-form deformations. IEEE transactions on medical imaging, 22(1):120–128, 2003.

Paganin, D., Mayo, S. C., Gureyev, T. E., Miller, P. R., and Wilkins, S. W. Simultaneous phase and amplitude extraction from a single defocused image of a homogeneous object. Journal of microscopy, 206(1):33–40, 2002.

Pichat, J., Iglesias, J. E., Yousry, T., Ourselin, S., and Modat, M. A survey of methods for 3D histology reconstruction. Medical image analysis, 46:73–105, 2018.

Rehberg, M., Krombach, F., Pohl, U., and Dietzel, S. Label-free 3d visualization of cellular and tissue structures in intact muscle with second and third harmonic generation microscopy. PloS one, 6(11):e28237, 2011.

Reiser, J., Albers, J., Svetlove, A., Mertiny, M., Kommoss, F. K., Schwab, C., Schneemann, A., Tromba, G., Wacker, I., Curticean, R. E., et al. Integrative imaging of lung micro structure: Amplifying classical histology by paraffin block µct and same-slide scanning electron microscopy. bioRxiv, pages 2024–06, 2024.

Sagar, M. M. R., Svetlove, A., D’Amico, L., Pinkert-Leetsch, D., Missbach-Guentner, J., Longo, E., Tromba, G., Bohnenberger, H., Alves, F., and Dullin, C. Optical clearing: an alternative sample preparation method for propagation based phase contrast µCT. Frontiers in Physics, 12:1433895, 2024.

Schindelin, J., Arganda-Carreras, I., Frise, E., Kaynig, V., Longair, M., Pietzsch, T., Preibisch, S., Rueden, C., Saalfeld, S., Schmid, B., et al. Fiji: an open-source platform for biological-image analysis. Nature methods, 9(7):676–682, 2012.

Song, Y., Treanor, D., Bulpitt, A. J., and Magee, D. R. 3D reconstruction of multiple stained histology images. Journal of pathology informatics, 4(2):7, 2013.

Sun, A. K., Fan, S., and Choi, S. W. Exploring Multiplex Immunohistochemistry (mIHC) techniques and histopathology image analysis: Current practice and potential for clinical incorporation. Cancer Medicine, 14(1):e70523, 2025.

Sung, H., Ferlay, J., Siegel, R. L., Laversanne, M., Soerjomataram, I., Jemal, A., and Bray, F. Global cancer statistics 2020: GLOBOCAN estimates of incidence and mortality world-wide for 36 cancers in 185 countries. CA: a cancer journal for clinicians, 71(3):209–249, 2021.

Svetlove, A., Griebel, T., Albers, J., D’Amico, L., Nolte, P., Tromba, G., Bohnenberger, H., Alves, F., and Dullin, C. X-ray phase-contrast 3D virtual histology characterises complex tissue architecture in colorectal cancer. Frontiers in Gastroenterology, 2:1283052, 2023.

Takeda, T., Momose, A., Hirano, K., Haraoka, S., Watanabe, T., and Itai, Y. Human carcinoma: early experience with phase-contrast x-ray ct with synchrotron radiation—comparative specimen study with optical microscopy. Radiology, 214(1):298–301, 2000.

Yakovlev, M. A., Vanselow, D. J., Ngu, M. S., Zaino, C. R., Katz, S. R., Ding, Y., Parkinson, D., Wang, S. Y., Ang, K. C., La Riviere, P., et al. A wide-field micro-computed tomography detector: micron resolution at half-centimetre scale. Synchrotron Radiation, 29(2):505– 514, 2022.

